# Transcriptome analysis of human preimplantation embryo reveals expressed waves associated with blastulation failure based on embryonic grade and age

**DOI:** 10.1101/2022.06.02.494565

**Authors:** Ping Yuan, Ying Liu, Haijing Zhao, Guangwei Ma, Lingyan Zheng, Qingxue Zhang, Hui Chen, Wenjun Wang, Yabin Guo

**Affiliations:** IVF Center, Department of Obstetrics and Gynecology, Sun Yat-sen Memorial Hospital, Sun Yat-sen University, Guangzhou, 510120, China; IVF Center, The First People’s Hospital of Kashi Prefecture, Kashi, 843800, China; Guangdong Provincial Key Laboratory of Malignant Tumor Epigenetics and Gene Regulation, Guangdong-Hong Kong Joint Laboratory for RNA Medicine, Medical Research Center, Sun Yat-sen Memorial Hospital, Sun Yat-sen University, Guangzhou, 510120, China; Ministry of Education Key Laboratory for Ecology of Tropical Islands, Key Laboratory of Tropical Animal and Plant Ecology of Hainan Province, College of Life Sciences, Hainan Normal University, Haikou, 571158, China

**Keywords:** Transcriptome, RNA sequencing, preimplantation embryo, blastulation failure, gene expression

## Abstract

In the *in vitro* fertilization and embryo transfer (IVF-ET) treatments, blastocyst culture is the method of choice for the generation of the embryos. Blastocysts can present different growth, quality, availability, and morphological characteristics that can be used to evaluate them. Although extreme blastocyst formation failures have been associated with the alteration of a single gene, the molecular factors responsible for arrested embryos remain unknown. RNA-sequencing (RNA-seq) is a promising tool for facilitating transcriptomic studies in early human embryos, thus allowing the investigation of gene expression discrepancies associated with different morphological criteria. Herein, we performed transcriptome analyses of the different stages of arrested human embryos. We identified candidate genes and related cell signaling pathways potentially associated with either arrested or developed embryos. Specifically, the three genes (*MOV10L1, DDX4*, and *FKBP6*) related to both DNA methylation and piRNA metabolic pathway might be involved in embryo development. Additionally, the transcriptome of arrested early blastocysts was significantly different from developed late blastocysts. Although the gene expression profiles identified were not significantly different between low- and high-quality late blastocysts, a significant difference in the profiles of day 5 and day 6 available late blastocysts was observed, which may be related to the clinical pregnancy rate associated with IVF-ET. Furthermore, we show that some chimeric RNAs may be functional in blastocyst development. Our findings uncovered new molecular markers that can be used for embryonic development detection, which might act as a tool for blastocyst selection for subsequent transfer.

## Introduction

During *in vitro* fertilization and embryo transfer (IVF-ET) treatments, blastocyst culture and single blastocyst transfer strategies have been used to select the most viable embryos to increase clinical pregnancy as well as overcome implantation failure (Gardner et al. 2000; Hardarson et al. 2012). Therefore, blastocyst quality during blastulation is an important indicator of the developmental potential of cleavage-stage embryos. However, approximately 50 % of normal fertilized cleavage embryos fail to form a viable blastocyst on day 6, as their development may get arrested at the cleavage, morula, early blastocyst, or unavailable late blastocyst stages even if they are cultured with high-quality cleavage embryos since day 3 (Gardner et al. 2000). This phenomenon is defined as embryonic developmental arrest (Niakan et al. 2012). However, the mechanisms of blastocyst formation failure and the reasons for the different developmental arrest stages in cleavage embryos remain unclear. Unexpectedly, embryos with less than seven cells, defined as being in the low-quality cleavage stage, succeeded to enter blastulation even in high-quality blastocysts, whereas some high-quality cleavage embryos scored as 8-cell grade 1 fail to enter the blastula stage (Yu et al. 2018). Whether the gene expression profiles of blastocysts differ among different cleavage embryos remains to be clarified.

Human preimplantation embryonic development includes a series of dynamic morphological changes from the formation of male and female pronuclei (two-pronuclear [2PN] zygote) to cleavage, compaction of morula, and eventually the formation of blastocyst that contains an outer trophectoderm (TE) and an inner cell mass (ICM) (Niakan et al. 2012; Xue et al. 2013). The two key morphological events that occur after the cleavage stage are compaction and cavitation. During blastocyst formation, compaction is characterized by intercellular tight interactions that form an early 16-cell morula to a late 32-cell morula after the 8-cell stage on day 3. Cavitation is the formation of a small single cavity within the embryo, which is a morphological marker of blastulation (Hardarson et al. 2012). Blastocyst quality is determined by the expansion of TE lineages and the cell number of ICM lineages (Niakan et al. 2012; Gardner et al. 1999). Previous studies have shown that the fate of TE lineages segregating from ICM lineages is determined by the GATA3/TEAD4/CDX2 axis in mice, and OCT4 (POU5F1) is required for continued repression of trophoblast fate both in mouse and human blastocysts (Ralston et al. 2010; Chazaud et al. 2016). Moreover, recent studies have analyzed some new molecular markers of TE and ICM in human blastocysts using single-cell RNA-sequencing (scRNA-seq), including GATA3/NR2F2 as TE markers and IFI16 as an ICM marker (Gerri et al. 2020; Meistermann et al. 2021). Whether these markers show differential expression in developmentally arrested embryos at different stages remains unclear. Another RNA-sequencing (RNA-seq) study of trophectoderm biopsy and the remaining whole embryo studies have revealed that poor-quality blastocysts are associated with oncogene activation and altered transcription of cell adhesion or extracellular matrix genes (Groff et al. 2019). Additionally, maternal mRNA decay defects and abnormal chromosomes are considered to be related to blastocyst developmental arrest (Qi et al. 2014; Sha et al. 2020). However, the underlying cause of blastocyst developmental arrest remains uncertain.

To address the above-mentioned gaps in knowledge, we used RNA-seq to analyze differently expressed genes related to blastulation failure, morphological grades, and embryonic age. We also provided a proof-of-principle dataset of whole-embryonic transcriptome data for identifying candidate genes associated with embryo development arrest and evaluating early human embryo development.

## Results

Sixty embryos from 21 couples that underwent preimplantation genetic testing (PGT) were included in this study. Among the 21 couples, 13 carried reciprocal translocations (REC), 2 carried inversions (INV), 2 carried Robertsonian translocations (ROB), 3 carried normal karyotyping in PGT for aneuploidies (PGT-A), and 1 carried normal karyotyping in PGT for monogenic diseases (PGT-M). The average age of women was 31.6 years and that of the men was 33.7 years. Sixty donated whole embryos were thawed and processed for whole-transcriptome amplification and RNA-seq analysis. To evaluate the transcriptome profiling in different developmental stages of human embryos, the embryo score groups were subdivided into five categories (for details, see Materials and Methods): cleavage (C, n = 18), early blastocyst (EB, n = 18), unavailable blastocyst (UB, n=5), low-quality blastocyst (LB, n = 10), and high-quality blastocyst (HB, n = 9) (Fig. 1B, Supplemental Table 1).

**Fig. 1.**
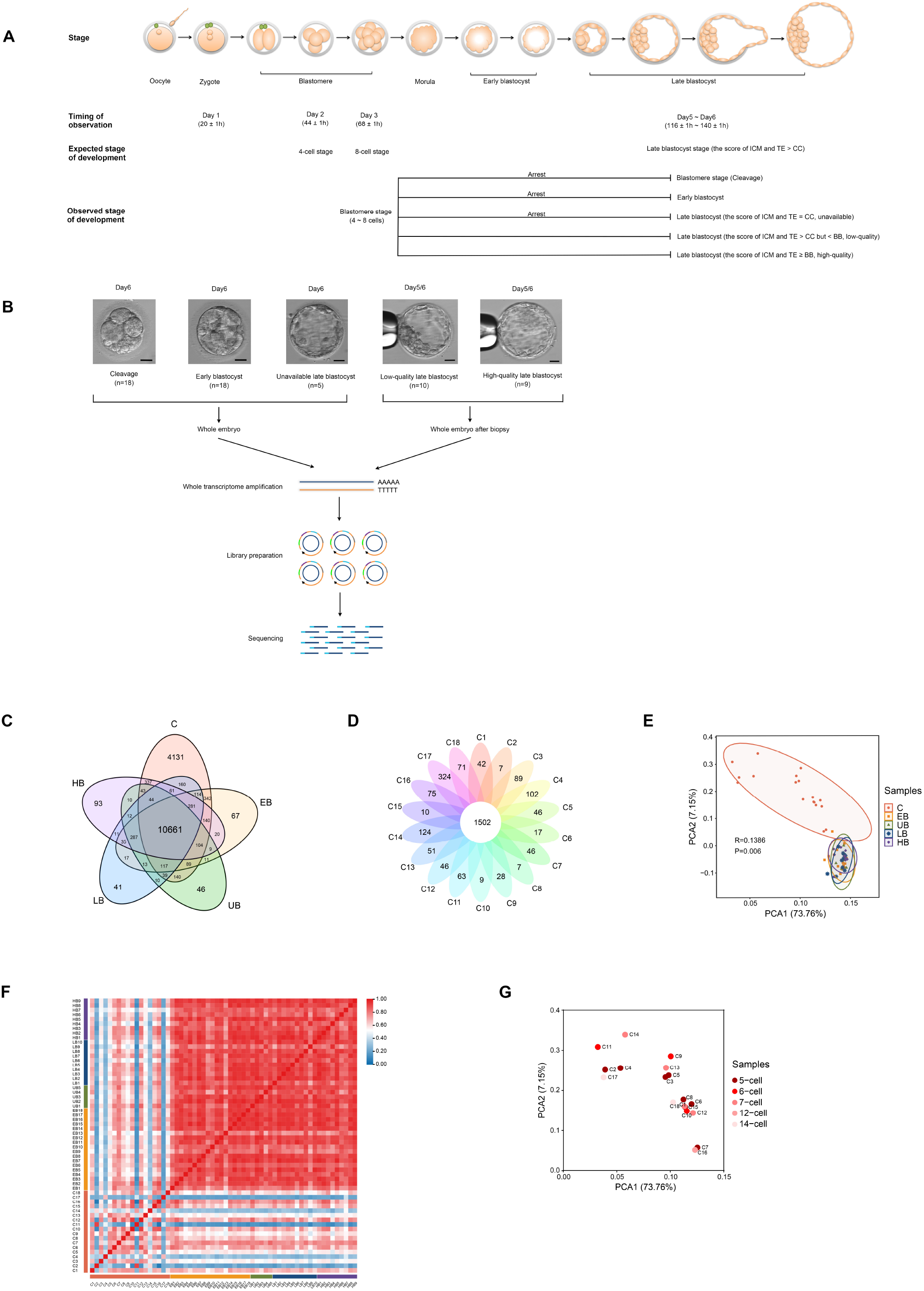
Expression levels of genes at different stages and correlation between samples. A) Schematic illustration of preimplantation embryo development. We highlighted the link between the observation time, expected developmental stages, and observed developmental stages for developmentally arrested blastocysts in the cleavage stage, early blastocyst stage, or unavailable blastocyst stage on day 6. B)Sixty embryos undergoing blastocyst culture were divided into five groups. Arrested embryos and available late blastocyst samples were processed. The arrested embryos were divided into three groups: cleavage, early blastocyst, and unavailable late blastocyst. Available late blastocysts were divided into low- and high-quality late blastocysts. Whole embryos or the remaining whole embryos after the preimplantation genetic testing biopsy were analyzed using RNA sequencing. Scale bars: 50 μm. C)Venn diagram of gene expression in each group. The number in the center of each circle represents the number of genes expressed in all groups (genes with an average transcripts per kilobase of exon model per million mapped reads (TPM) ≥ 1 in all groups were counted), and the number in each petal represents the genes specifically expressed in each group. D)Petals diagram of gene expression in cleavage embryos. The numbers in the central circles indicate the number of genes expressed in 18 embryos at the cleavage stage (genes with a TPM ≥ 1 in all samples were counted), and the number in each petal represents the genes specifically expressed in each sample. E)Principal component analysis (PCA) of total samples. The X- and Y-axes indicate the corresponding principal components obtained after dimensionality reduction of the sample expression. P represents significance, and R represents the goodness of fit index. F) Correlation heatmap of all samples. The color bar indicates the Pearson’s correlation coefficient. G) PCA of samples at the cleavage stage.

### Transcriptional profiles across the five different groups

In total, we generated 472.18 Gb of raw data corresponding to the transcriptome profiles of 60 whole embryos. The average clean data output for each sample was 6.58 Gb (Supplemental Table 4). Then 394.95 Gb (clean data) of filtered reads were processed for quality control (QC, Supplementary Table 2). A total of 235.84 Gb of filtered reads, 30.5% of initial data, were mapped to the RefSeq human genome, and an average of 4.27% of data were mapped to the mitochondrial genome (Supplemental Table 4, Supplemental Fig. 1A). The analysis identified 19,764 out of 20,040 genes with TPM ≥ 1 in at least one sample (Supplemental Fig. 1B). First, we analyzed the number of known genes expressed in each of the 60 embryos. On average, we detected the expression of 10,661 (53.20%) out of 20,040 RefSeq genes (genes with TPM ≥ 1 in every sample were counted), which were expressed in all groups (Fig. 1C, Supplemental Table 5). More than half of the known human genes were expressed in the sampled embryos. Second, we compared the gene expression in each of the five groups. The percentage of genes with TPM < 10 in group C was significantly higher than that in the other groups (*P* = 0.002). With the orderly expression of genes associated with embryonic development, highly expressed genes (TPM ≥ 10) were maintained at a stable ratio in blastocysts (Supplemental Fig. 1B). As group C was characterized by 4,131 uniquely expressed genes, we evaluated the gene expression levels of 18 samples in group C and discovered that only 1,502 genes (7.60%, 1502/19,764) were expressed in all the samples (Fig. 1D), indicating the great diversity among the expression profiles of the embryos in group C.

To determine whether these gene expression profiles correlated with different developmental stages, we analyzed the RNA-seq data of embryos in the blastomere and blastocyst stages. The greatest changes in gene expression were observed between the blastomere and blastocyst stages, and gene expression at different embryonic stages had specific clusters, which were highlighted by principal component analysis (PCA; Fig. 1E). However, there were no remarkable differences among the other four groups at the blastocyst stage, which indicated that cavitation formed with a single cavity through fluid pumping should be a pivotal morphological event. To determine the correlation of gene expression between the experimental groups, we calculated the Pearson correlation coefficient between each sample pair, which revealed the apparent transcriptome difference between the blastomere and blastocyst stages. However, the transcriptome profiles of 42 blastocysts were relatively similar (Fig. 1F), and there also were no differences among 18 arrested cleavage-stage embryos (Fig. 1G).

From the perspective of embryo development and blastocyst formation, tight interactions and single cavities gradually formed between cells after 8-cell cleavage. Almost all high-quality cleavage embryos (score on day 3 was 8-cell grade 1) skipped the arrest at the morula stage and progressed to the blastocyst stage (blastocyst formation rate of 97.30% (36/37)), except for one that arrested at the 14-cell stage (Supplemental Table 1). However, among another 23 embryos containing 4–7 cells on day 3, only 6 proceeded to the blastocyst stage, most of which stayed at a standstill except for 2 arrested at the 12- and 14-cell stages (Supplemental Table 1). This indicated that the formation of compaction and cavitation would be difficult to overcome for 4- to 7-cell cleavage embryos, but not for high-quality 8-cell cleavage embryos.

### Embryo grades exhibit differential expression

We first compared the differences in gene expression among the five developmental arrest stages of the preimplantation embryos. The greatest changes in gene expression were observed between groups C and EB, while there were no significant differences between the LB and HB groups (Fig. 2A-2D, Supplemental Fig. 2A). The number of differentially expressed genes (DEGs) decreased gradually among the four groups as the embryo development progressed (Fig. 2A-2D, Supplemental Fig. 2A). First, *ACTL8* (Q = 6.44 × 10^-36^) and *EIF4E1B* (Q = 9.11 × 10^-32^) were two of the genes significantly upregulated in group C than in group EB (Fig. 2A). ACTL8 is an actin-like protein that may act as a tumor facilitator (Yao et al. 2014; Li et al. 2020). EIF4E1B is associated with ribosome assembly in the majority of eukaryotic mRNAs (Kropiwnicka et al. 2015). Additionally, *HNRNPCL1* (Q = 5.05 × 10^-20^) and *GCOM1* (Q = 1.08 × 10^-13^), which are also involved in cancer, were highly upregulated in group EB and group UB respetively (Fig. 2B-2C) (Schinke et al. 2014; Gao et al. 2020). Together, these data suggest that preimplantation embryos with developmental arrest showed a lack of transcriptional control reminiscent of the hallmarks of cancer, and embryos arrested at the cleavage stage might be associated with unfulfilled mRNAs.

**Fig. 2.**
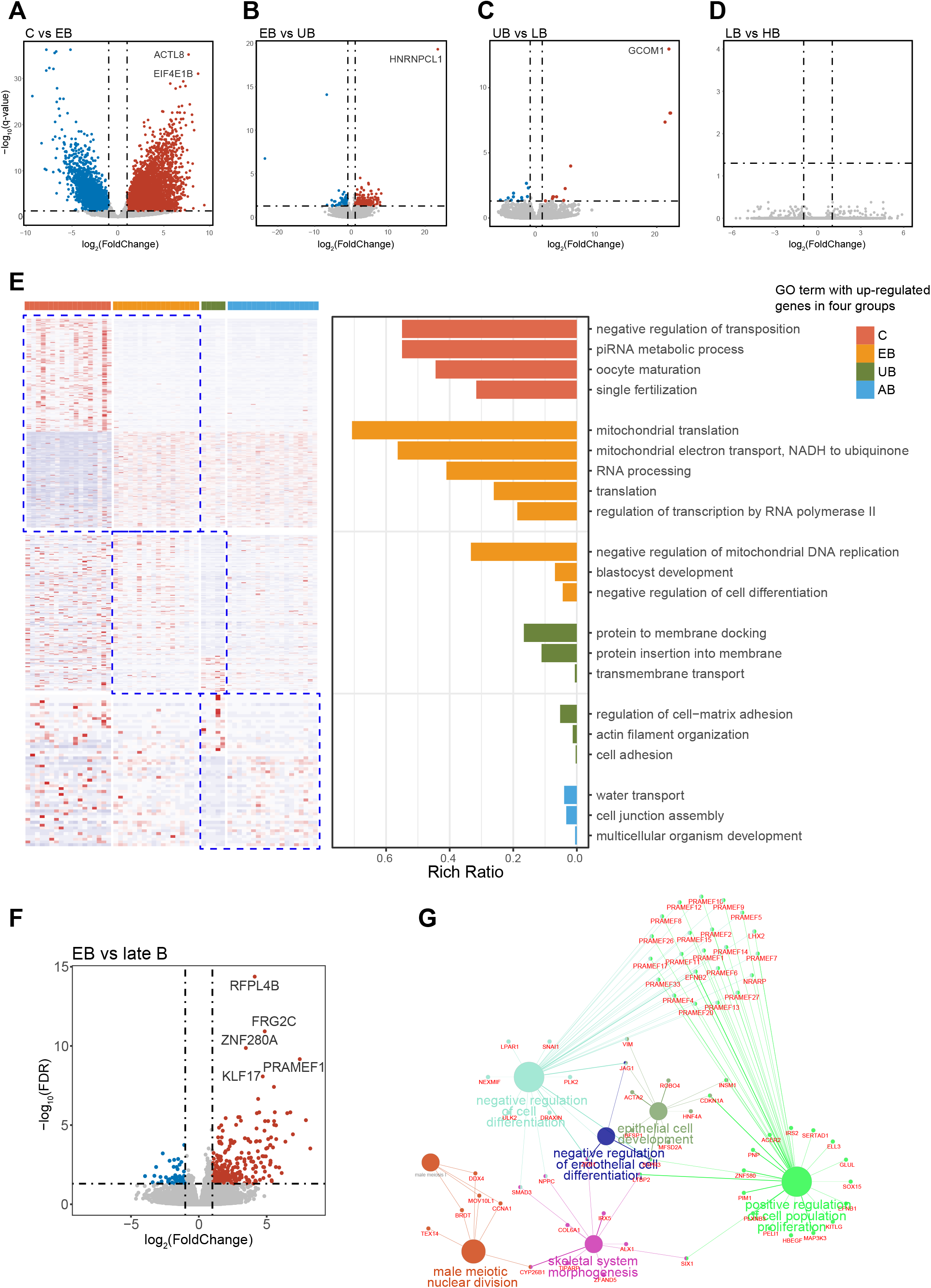
Differentially expressed gene (DEG) and enrichment analyses. A-D) Volcano plots. DEGs in cleavage embryos vs early blastocysts (A), early blastocysts vs unavailable blastocysts (B), unavailable blastocysts vs low-quality blastocysts (C), and low-quality blastocysts vs high-quality blastocysts (D). E) Heatmap of DEGs between adjacent stages and Gene Ontology (GO) enrichment analysis. F) Volcano plot of DEGs in cleavage embryos vs late blastocysts. G) GO enrichment network of upregulated genes in early vs late blastocysts.

To gain insight into the potential functional differences of the dynamically changing genes in different developmental arrest stages, we subdivided the 60 embryos into four groups: group C, group EB, group UB, and group AB (group AB represented available late blastocysts comprised from both LB and HB groups). We performed a Gene Ontology (GO) analysis on the DEGs and found that the enrichment of GO terms was different in each comparison group. Comparing groups C and EB, upregulated genes in the cleavage arrest stage were found to be enriched for the functional categories of negative regulation of transposition, piRNA metabolic processes, oocyte maturation, and single fertilization (Fig. 2E). However, in the early blastocyst arrest stage, upregulated genes were mainly enriched for mitochondrial functions, especially nicotinamide adenine dinucleotide (NADH), RNA processing, and translation (Fig. 2E), thus indicating that embryos arrested at the cleavage stage might be involved in abnormal metabolism of piRNA and NADH-related mitochondria (Russell et al. 2017; Santos Monteiro et al. 2021). The embryos in the UB group exhibited enlarged cavitation through fluid pumping compared to those in the EB group, which displayed enrichment of protein to membrane docking pathway as well as protein insertion into the membrane pathway (Fig. 2E). This implied that the TE was formed, and the water flow in and out of the blastocoel was established in late blastocysts, despite embryos being arrested in the unavailable late blastocysts. However, the negative regulation of cell differentiation and mitochondrial DNA replication also affected the development from EB to UB. When comparing groups UB and AB, the AB group showed improved water transport, multicellular organism development, and cell junction assembly (Fig. 2E), indicating that better transportation and cell junction assembly might increase the cell number and the expansion of the blastocoel.

To investigate the early blastocyst cell fate decision and differentiation of TE and ICM, we compared the EB group with the late B group (group UB + group LB + group HB). Cell proliferation as well as the differentiation of TE and ICM were the major morphological differences between late and early blastocysts. We identified 334 DEGs between the EB and B groups (Fig. 2F). The five most significantly upregulated genes in the EB group included *RFPL4B, FRG2C, ZNF280A, PRAMEF1*, and *KLF17* (Fig. 2F, Supplemental Fig. 3). Interestingly, *ZNF280A*, a transcription factor, inactivates the Hippo signaling pathway-dependent proliferation and tumorigenesis in colorectal cancer, as the Hippo pathway plays a vital role in the initiation of TE fate in human embryo development (Wang et al. 2019; Gerri et al. 2020). PRAMEF1 is also a potential biomarker in cancer, and its upregulation may be associated with a lack of transcriptional control (Gao et al. 2020). KLF17, a transcription repressor, is enriched in 8-cell, morula, as well as early blastocyst stages and might be associated with maternal genes in human embryos (Meistermann et al. 2021). *RFPL4B* and *FRG2C* (DUX4-activated genes) were significantly higher in the EB group, which implies that embryonic genome activation (EGA) is involved in the early blastocyst arrest stage (Hendrickson et al. 2017). To identify the pathways associated with the 334 DEGs, we performed a GO analysis which revealed that significantly upregulated genes were associated with positive regulation of cell proliferation and negative regulation of cell differentiation (Fig. 2G). As shown in Fig. 2G, almost all members of the PRAME gene family appeared in the significantly upregulated gene set (Fig. 2G). The leucine-rich repeat proteins encoded by the PRAME gene family function as transcription regulators in cancer cells and also play important roles in spermatogenesis and oogenesis (Birtle et al. 2013). Together, these data suggest that the arrest of cell proliferation might be associated with enrichment of the KLF17-related transcription repressor and the members of the PRAME gene family, while ZNF280A, a key transcription factor capable of inactivating the Hippo pathway, would influence the TE differentiation in human early blastocysts.

To assess blastocyst development from different quality cleavage embryos on day 3, we divided embryos into two subgroups: 4C_AB and 8C_AB (Supplemental Table 1). 4C_AB represents available late blastocysts derived from 4 to 6 cells cleavage embryos, and 8C_AB represents available late blastocyst derived from 8 cells cleavage embryos. Our results revealed six significantly upregulated genes in group 4C_AB (Supplemental Fig. 2B-2C). TRDMT1 is believed to be responsible for the methylation of aspartic acid transfer RNA, increased tRNA stability, and steady-state protein synthesis (Tuorto et al. 2012). PYROXD2, a probable oxidoreductase, may regulate mitochondrial function (Wang et al. 2019). Collectively, these data indicate that morphologically low-quality cleavage embryos on day 3, which eventually formed available late blastocysts, might be capable of activating gene expression in response to protein synthesis and mitochondrial function required at the blastocyst stage.

### Dynamic patterns of embryo gene expression during different arrest stages

To investigate the dynamic gene expression changes during the different arrest stages of preimplantation embryos, we subdivided the embryos into three groups: group C, group EB, and group late B. We discovered that 50 genes were successively upregulated and 11 genes were successively downregulated in the order C-EB-late B (Fig. 3A). Furthermore, we performed a GO analysis on the upregulated DEGs in group C compared them with those in the other two groups and found that the most significantly enriched GO terms were associated with DNA methylation involved in gamete generation and Piwi-interacting RNAs (piRNA) metabolic processes, of which three genes (*MOV10L1, DDX4*, and *FKBP6*) were shared in both pathways (Fig. 3B).

**Fig. 3.**
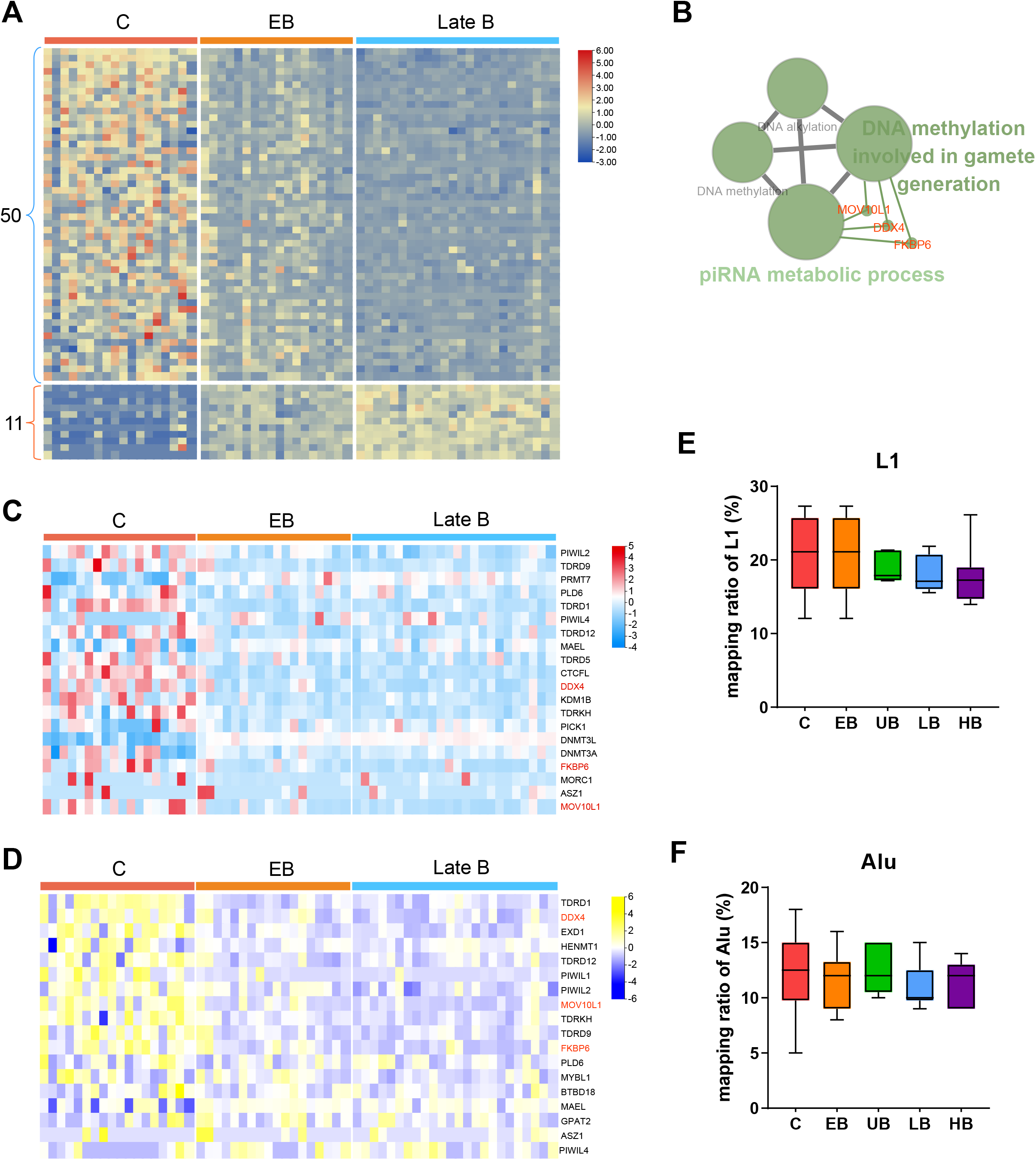
Gene expression model of embryos at different stages. A) Heat map of two sets of genes. There were 50 genes that were characterized by gradually decreasing expression levels from the cleavage stage to late blastocyst stage (log_2_FC (C vs. EB) > 1 and log_2_FC (EB vs. LB) > 1), while the expression of 11 genes increased (log_2_FC (C vs. EB) < -1 and log_2_FC (EB vs. LB) < -1). The row indicates the gene and the column indicates the sample. B)Gene Ontology (GO) enrichment analysis of upregulated differentially expressed genes (DEGs) in the cleavage stage (case group) vs other stages (control group), performed using ClueGO. The size of the nodes indicates their significance and the most significant pathways are illustrated with the largest node size. The red front shows the genes shared between the two terms. C) Expression heatmap of genes related to DNA methylation involved in gamete generation. The red font indicates the genes shared between the two pathways in Fig. 3B. D) Expression heatmap of genes related to piRNA metabolic processes. The genes given in red font are the genes shared between the two pathways in Fig. 3B. E-F) Mapping ratios of L1 (E) and Alu (F). Black lines indicate median values and boxes range from the 25^th^ to 75^th^ percentiles. The whiskers indicate the ratio from max to mix. FC: fold change; C: cleavage; EB: early blastocyst; LB: low-quality blastocyst.

PiRNAs are involved in the post-transcriptional gene silencing of transposable elements through the slicer activity of piRNA-induced silencing complexes (piRISCs) (Russell et al. 2017). Some piRNAs also appear to participate in transcriptional gene silencing, primarily by directing DNA methylation and histone modifications (Carmell et al. 2007; Klenov et al. 2014). We examined the expression levels of the genes related to both DNA methylation and piRNA metabolic processes and found that most of these genes were highly expressed at the cleavage stage (Fig. 3C and 3D), indicating that the arrested cleavage embryos might not have successfully suppressed the piRNA pathway, resulting in the failure of the embryo genome expression reprogramming process.

While transposable element transposition is often associated with negative outcomes and is recognized as a threat to the host genome, the expression of LINE-1 (L1), the most abundant transposable element in the human genome, appears to be required as an integral part of the transcriptional networks regulating cellular potency during the early development of mammalian embryos (Percharde et al. 2018). The transcripts of SINEs, LINEs, and long terminal repeat (LTR) retrotransposons were relatively abundant before the 8-cell stage; they then gradually decreased to the basal level in post-implantation embryos (Guo et al. 2014). However, the gene expression levels of L1 and Alu were not significantly different among the five blastocyst groups (p = 0.2094 for L1 and p = 0.6351 for Alu; Fig. 3E and 3F).

### Embryo age influences differential gene expression

Within the AB group, no significant DEGs were observed between the LB and HB groups (Fig. 2D). To determine the difference in gene expression in late available blastocysts, we performed a hierarchical cluster (H-cluster), analysis and the late blastocysts were classified into two major clusters (Fig. 4A). The embryos in the left cluster were B5s and those in the right cluster were B6s. Gene expression distributions differed between the B5 and B6 groups (Fig. 4A), with a total of 255 genes being differentially expressed (Supplemental Fig. 4). *KRT19* (Q = 6.199 × 10^-10^) was more significantly upregulated in group B6 than in group B5 (Supplemental Fig. 4). KRT19 is a type I cytokeratin specifically expressed in the periderm or developing epidermis and may be involved in cell differentiation (Gupta et al. 2017). Furthermore, we performed GO analysis for these DEGs and discovered that the upregulated genes in group B5 were enriched for the functional categories of positive regulation of cell migration and gene expression, whereas the upregulated genes in group B6 were enriched for the functional categories of negative regulation of the timing of anagen and cellular response to growth factor stimulus, negative regulation of cell-matrix adhesion, cell adhesion mediated by integrin, DNA damage response, and signal transduction by p53 class mediator, thus resulting in cell cycle arrest (Fig. 4B).

**Fig. 4.**
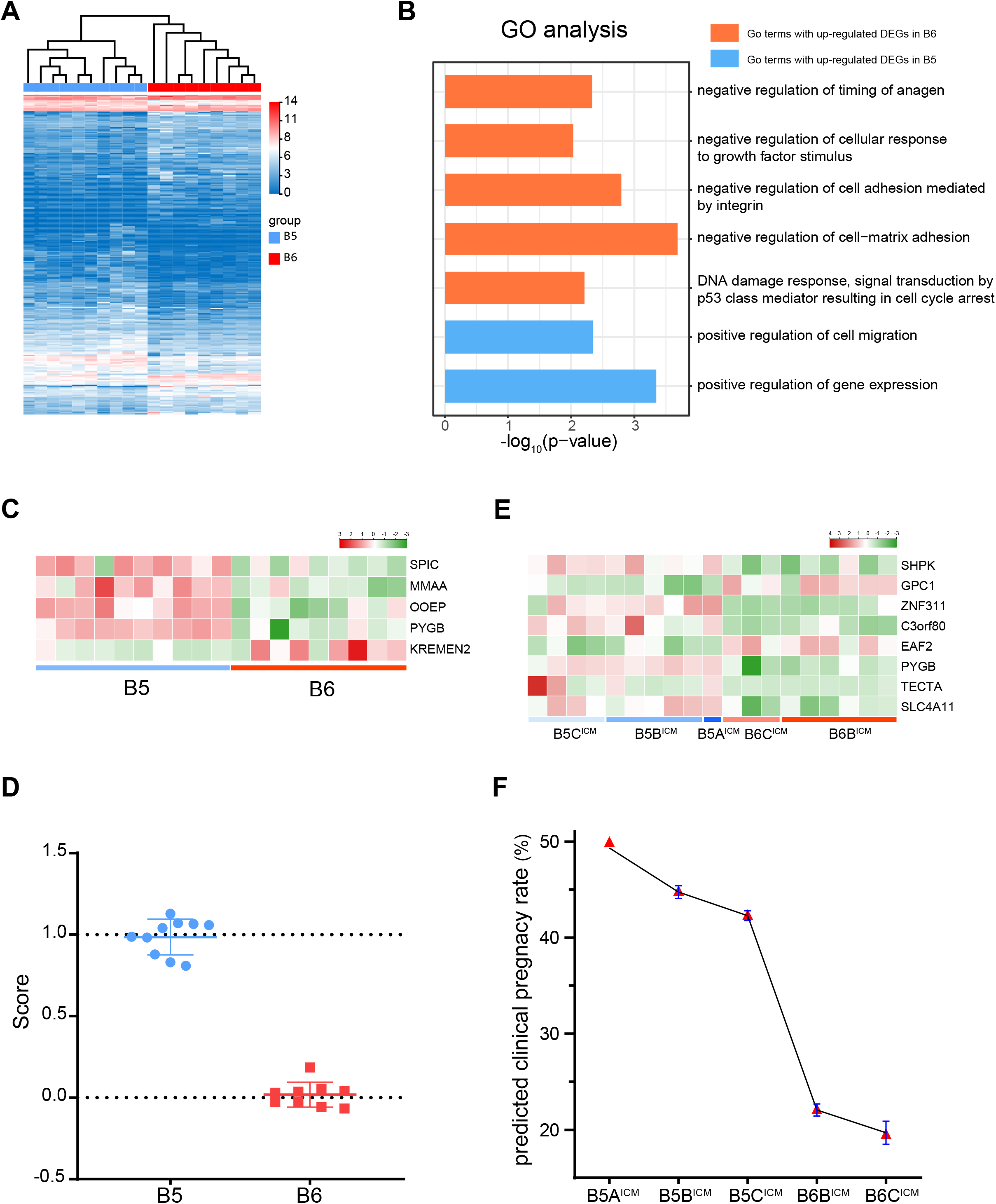
Gene expression difference in available late blastocysts. A)Hierarchical (H)-cluster of total gene expression of available late blastocysts. B)Gene Ontology (GO) enrichment analysis of differentially expressed genes (DEGs) between 5Bs and 6Bs. C)Heat map of gene expression levels in the available blastocyst fitting equation. D)Heat map of gene expression levels in the clinical pregnancy rate (CPR) fitting equation. E)The available blastocyst score distribution was calculated according to the fitting equation (mean ± SD). F)CPR predicted by the fitting equation (mean ± SD in blue). The red triangle represents the CPR corresponding to the embryo grade according to the clinical data from the Sun Yat-sen Memorial Hospital Reproduction Center from 2011 to 2018.

To define a gene signature able to distinguish groups B5 and B6 based on RNA profiles, we selected five DEGs (*SPIC, MMAA, OOEP, PYGB*, and *KREMEN2*, Fig. 4C), and calculated the linear regression using the following formula: embryo age = -0.282 + 0.163 × MMAA - 0.036 × KREMEN2 + 0.28 × SPIC + 0.005 × OOEP + 0.003 × PYGB (Supplemental Table 2). Using this formula, the scores of group B6 were close to 0, whereas those of group B5 were close to 1. This score difference between the groups was significant and there was no overlap (Fig. 4D).

Our data also revealed that the clinical pregnancy rate (CPR) was higher in group B5 than in group B6 (45.5% (173/380) vs. 21.9% (33/151), P = 4.419 × 10^-7^). Next, to assess the CPR associated with different ICM grades of available late blastocysts, we divided blastocysts into five subgroups: B5A^ICM^, B5B^ICM^, B5C^ICM^, B6B^ICM^, and B6C^ICM^ (Supplemental Table 1). B5A^ICM^ represents group B5 with ICM graded as A; group B5B^ICM^ represents group B5 with ICM graded as B; group B5C^ICM^ represents group B5 with ICM graded as C; group B6B^ICM^ represents group B6 with ICM graded as B; group B6C^ICM^ represents group B6 with ICM graded as C. According to the five subgroups, we identified eight DEGs (*TECTA, EAF2, ZNF311, GPC1, C3orf80, PYGB, SHPK*, and *SLC4A11*; Fig. 4E), and derived the CPR formula using the different blastocyst scores: CPR = 13.344 + 2.616 × TECTA + 1.632 × EAF2 + 1.442 × ZNF311 - 1.132 × GPC1 + 1.108 × C3orf80 + 0.219 × PYGB -0.145 × SHPK - 0.01 × SLC4A11 (Supplemental Table 3). The CPRs were found to gradually decrease for the five subgroups (B5A^ICM^, B5B^ICM^, B5C^ICM^, B6B^ICM^, and B6C^ICM^) with the corresponding values being 50.0% (20/40), 44.9% (105/234), 42.4% (28/66), 33.3% (2/6), 22.2% (22/99), and 19.6% (9/46) (Fig. 4F). This implies that the expression levels of TECTA, EAF2, ZNF311, GPC1, C3orf80, PYGB, SHPK, and SLC4A11 may be closely associated with CPR in blastocysts.

### Dynamics expression of chimeric RNAs

Chimeric RNAs and their encoded products are usually one of the causes of tumors (Wang et al. 2021) and our arrested embryos showed a lack of transcriptional control of cancer. To investigate chimeric RNAs associated with blastocyst development failure, we observed significantly different expression spectra of chimeric RNAs in different blastocyst groups; the chimeric RNAs in group C showed the greatest variance (Fig. 5A). Using EricScript analysis, we found a large number of chimeric RNAs in 60 samples, and each group had more than 10% chimeric RNAs, especially in groups C and EB (Fig. 5B-5F). Here, we termed the chimeric RNAs appearing in all samples of one group as common chimeric RNAs. In group C, the number of common chimeric RNA was 0, whereas there were 1–10 common chimeric RNAs in the other groups. We noticed that ZNF780B-LOC105372798 was detected in all samples of late blastocysts and most samples of early blastocysts (17/18), whereas it only appeared once in cleavage embryos (1/18) (Fig. 5G). In groups EB, UB, LB, and HB, the expression level of the ZNF780B-LOC105372798 chimeric transcript was higher than that of LOC105372798, but lower than that of ZNF780B (Fig. 5H). This indicates that some chimeric RNAs may be functional in early embryo development.

**Fig. 5.**
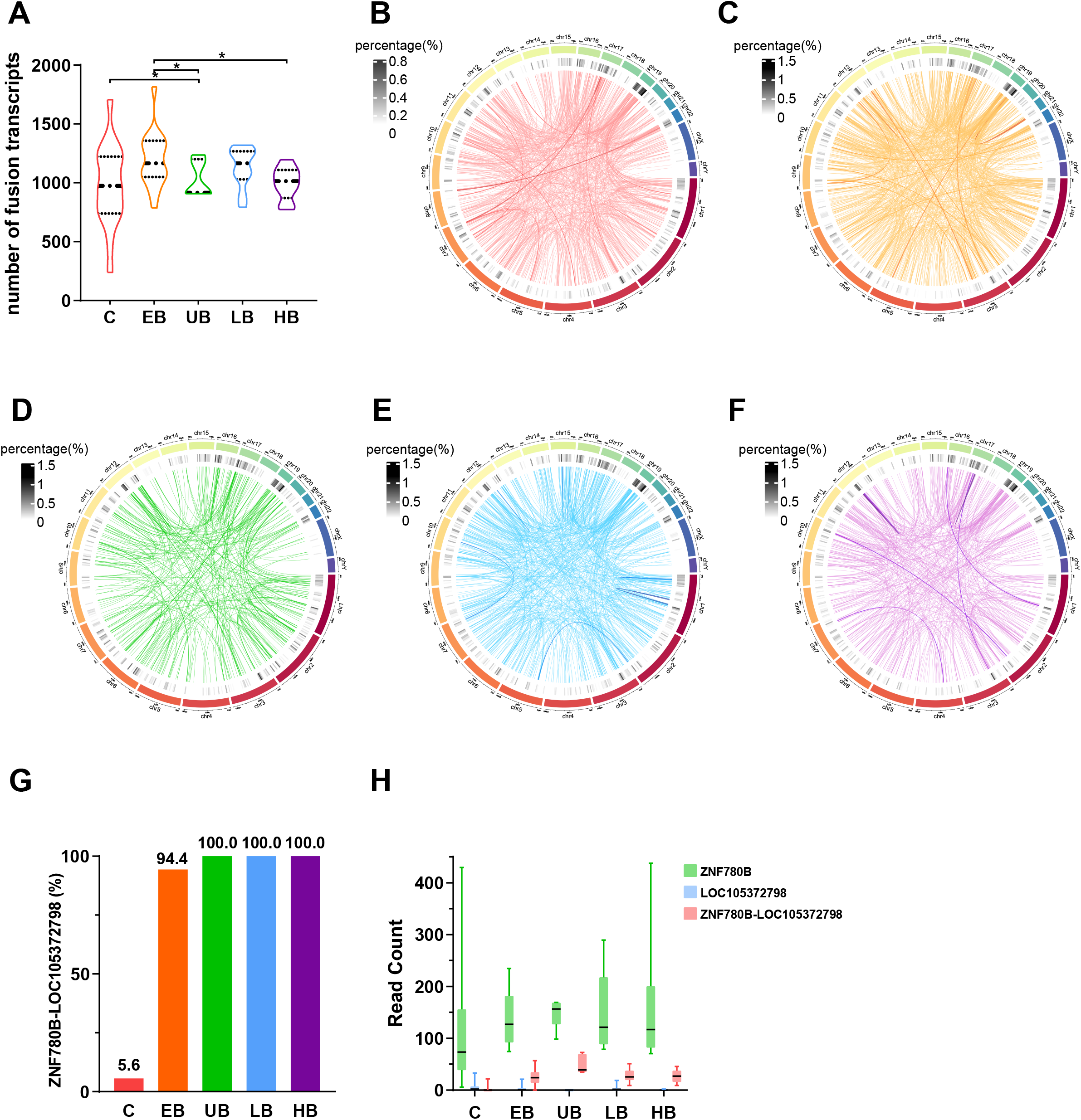
Chimeric RNAs from different stage embryos. A) Number of fusion RNAs. * indicates a significant difference between the two groups (p < 0.05). The dotted lines indicate the 25^th^ and 75^th^ percentiles and the median. The whiskers indicate the ratio from max to mix. Groups C and EB showed a greater variance in the number of fusion transcripts. B-F) Circos plots of chimeric RNAs in groups C (panel B), EB (panel C), UB (panel D), LB (panel E), and HB (panel F). The tracks of the Circos charts from the center to the outer part represent chimeric gene sites and the percentage of chimeric RNA in 5 Mbp sliding blocks. Links indicate the two genomic loci of chimeric RNAs. The color of the link indicates the frequency of chimeric RNA. Random 10% chimeric RNAs of total chimeric RNAs are shown in each plot. The legend is for the 1^st^ track, indicating the percentage of chimeric RNA in 5 Mbp sliding blocks. G) Frequency of chimeric RNA ZNF780B-LOC105372798 in each group. H) Read count of the chimeric transcript ZNF780B-LOC105372798 and read count of its two paternal genes, ZNF780B and LOC105372798. C: cleavage; EB: early blastocyst; UB: unavailable blastocyst; LB: low-quality blastocyst; HB: high-quality blastocyst.

## Discussion

Embryo development arrest during the preimplantation phase may result in a variety of morphological abnormalities. In this study, we identified developmental competence gene candidates and related cell signaling pathways obtained via transcriptomic analyses of arrested embryos. Specifically, the three genes (*MOV10L1, DDX4*, and *FKBP6*) related to both DNA methylation and piRNA metabolic pathway might be involved in blastocyst development. We found the transcriptome of arrested early blastocysts was significantly different from developed late blastocysts. Although the gene expression profiles identified were not significantly different between low- and high-quality late blastocysts, a significant difference in the profiles of day 5 and day 6 available late blastocysts was observed, which may be related to the clinical pregnancy rate associated with IVF-ET.

In this study, we painted a transcriptional picture of embryos under blastocyst culture in different developmental states, including information on samples from arrested whole embryos and available late blastocysts with chromosomal abnormalities. Previous studies have suggested that human embryos undergoing blastocyst developmental arrest are more likely to exhibit chromosomal abnormalities (Qi et al. 2014; Handyside et al. 2021). However, in a study by Fragouli et al, no detectable chromosomal aberration effects on morphology were detected at any preimplantation stage when analyzing the most clinically relevant aneuploidies (Fragouli et al. 2014). Our data confirmed that no significant differences in the blastulation rates between the PGT-SR group (preimplantation genetic testing for chromosomal structural rearrangement in couples) and the IVF group (traditional IVF couples with normal chromosol karyotypes) were observed by retrospective analysis from January 2019 to December 2019 (Supplemental Table 6). Conversely, the rates of available blastocysts in the PGT-SR group were significantly higher than those in the IVF group (Supplemental Table 6), implying that chromosomal abnormalities might not hinder preimplantation embryo development and blastulation.

In previous studies, single-cell transcriptome analysis using scRNA-seq revealed key fate markers in ICM cells, including epiblast (EPI), primitive endoderm (PrE), and TE cells during normal blastocyst development (Yan et al. 2013; Petropoulos et al. 2016; Gerri et al. 2020). However, the markers enriched in TE, EPI, and PrE were different. Markers with consistently high expression in TE were GATA3 and GATA2 while NANOG was associated with EPI and GATA4 was associated with PrE. Subsequent studies have integrated pseudotime analysis of the aforementioned scRNA-seq datasets and discovered that the fate markers IFI16 and GATA4 were specifically restricted to EPI and PrE (Meistermann et al. 2021). In our study, the expression of known markers was re-analyzed using RNA-seq of whole embryos with morphological differences. GATA4 was confirmed to be restricted in available late blastocysts, which implies that this marker can be used for the identification of ICM cells (Supplemental Fig. 3). However, in our data, IFI16 and NANOG markers were not only expressed in available late blastocysts but also in early blastocysts and cleavage embryos, suggesting that these markers might not be specifically expressed in EPI (Supplemental Fig. 3). Moreover, DNMT3L and GATA2 modules are broadly expressed in blastocysts but not in arrested cleavage embryos, which is consistent with the integrated pseudotime analysis (Supplemental Fig. 3; Meistermann et al. 2021). Additionally, the expression of multiple cancer-related markers (Fig. 2) was significantly upregulated in arrested embryos with poor morphological quality, which is in line with findings from a previous study (Groff et al. 2019). These upregulated tumorigenic markers might induce unordered transcriptional control and promote negative proliferation by inactivating the Hippo signaling pathway to cause embryonic developmental arrest (Wang et al. 2019). During maternal-to-zygotic transition (MZT) following fertilization, appropriate maternal RNA degradation and zygotic genome activation (ZGA) proceed in order. Previous studies have shown that ZGA is initiated on day 3 in human 8-cell embryos when abundant maternal RNA degradation occurs (Dobson et al. 2004; Vassena et al. 2011; Sha et al. 2020). A recent study demonstrated that human ZGA initiates at the 1-cell stage following fertilization, wherein the maternal RNA degradation begins (Asami et al. 2022). Therefore, maternal RNA degradation is believed to represent a vital process in embryonic development. Recently, a correlation between maternal RNA decay defects and early developmental arrest has been reported (Sha et al. 2020). Furthermore, the scRNA-seq results of the study from He et al showed that the expression of maternal RNA in the 8-cell stage embryos that underwent developmental failure in the future was higher than that in the same period of embryos undergoing successful blastocyst development (He et al. 2022). However, this study ignored the fact that cell number, cell size, and fragmentation also affect embryo development and blastulation (Alpha Scientists in Reproductive Medicine and ESHRE Special Interest Group of Embryology. 2011; Racowsky et al. 2011), and low-quality 8-cell embryos may fail in blastulation. Single cells from different-grade 8-cell embryos presented different gene expression patterns. Here, we compared the expression of some maternal genes, including a single gene associated with embryonic developmental arrest (Wang et al. 2018; Zheng et al. 2021a; Zheng et al. 2021b) between groups C and B (Supplemental Fig. 5). In accordance with a previous study (Yan et al. 2013), we show that the expression of maternal genes is significantly higher in the cleavage stage than in the blastocyst stage, whether by whole-embryo RNA-seq or scRNA-seq (Supplemental Fig. 5), which implies that failure of maternal RNA clearance may be an observable molecular manifestation in arrested embryos. Maternal RNA decay might be regulated by β-tubulin, the subcortical maternal complex, or other unknown factors (Zhu et al. 2021). Further studies are needed to elucidate the pathogenic mechanisms of maternal RNA clearance and ZGA associated with embryonic development.

Our GO analysis of upregulated DEGs showed that three genes (*MOV10L1, DDX4*, and *FKBP6*) related to both DNA methylation and piRNA metabolic pathway might affect embryo development arrest by suppressing embryo genome expression (Fig. 2E and Fig. 3). MOV10L1 as a germ cell–specific putative RNA helicase is associated with Piwi proteins, and *Mov10l1*^−/−^ male mice showed sterility and lack of piRNA (Zheng et al. 2010). *DDX4* encoding a DEAD box protein can mediate the repression of transposable elements and govern the methylation during meiosis via the piRNA metabolic process to be involved in spermatogenesis, and cellular growth and division (Castrillon et al. 2000; Li et al. 2010). Defects in *Fkbp6* gene in mice showed male infertility, indicating that FKBP6 as a component of the synaptonemal complex was essential for homologous chromosome pairing in meiosis (Crackower et al. 2003). A recent study also revealed that small RNA-mediated degradation of maternal transcripts influences cell division in embryos (Paloviita et al. 2021). It will be interesting to further investigate the association between DNA methylation and piRNA metabolic pathway and embryonic development.

Chimeric RNAs are generally considered to be the products of gene fusion or trans-splicing, implying active transcription issues. Our study showed that the frequencies of chimeric RNAs were higher in blastocysts than in cleavage embryos, especially for the fusion gene ZNF780B-LOC105372798 (Fig. 5). Notably, all chimeric points occurred at chr19:40032625-chr21:37561176, indicating that this chimeric RNA is not transcriptionally noisy, while a functional example of trans-splicing is transiently expressed at a specific developmental stage. ZNF780B-LOC105372798 is located between the exon region of ZNF780B and the exon region of a long non-coding RNA, which is more likely to be caused by changes in chromatin accessibility during reprogramming of early development, which supports the hypothesis that ZNF780B protein may be involved in transcriptional regulation (Tycko et al. 2020). It has been previously reported that the interference of some fusion RNAs could cause most cleavage embryos to fail in blastulation (Mao et al. 2018). Collectively, the increased transcripts in blastocysts imply EGA, which also suggests that chimeric RNAs play a vital role in early embryonic development.

In addition, the comparison of groups 4C_AB and 8C_AB revealed that more active mitochondrial function might promote the low-quality cleavage embryos to undergo blastocyst development by accelerating protein synthesis. (Supplemental Fig. 2B-2C). However, no DEGs were identified between the LB and HB groups (Fig. 2D and Supplemental Fig. 2A), which suggests that the slight morphological differences in available late blastocysts might not affect gene expression when the number of cells in blastocysts has reached a certain level.

Nonetheless, blastocysts of different ages exhibit differential gene expression (Petropoulos et al. 2016). Our study also described the differences between groups B5 and B6 in positive regulation of cell migration and gene expression, as well as the negative regulation of cell-matrix adhesion (Fig. 4). Furthermore, the key markers associated with CPR were identified to be highly expressed in group B5, but not in group B6 (Fig. 4). Together, these indicated that blastocysts on day 5 might exhibit a greater capacity for cell proliferation and adhesion compared to blastocysts on day 6, which is consistent with the fact that B5 embryos have better pregnancy outcomes (Haas et al. 2016; Bourdon et al. 2019).

Our RNA-seq analysis of whole embryos provides a complete expression profile while excluding the mosaicism effect from single cells or biopsies, which could be confounded by multiple expression signals due to different differentiated cells, especially at the blastocyst stage. Another limitation of this study was that the expressions of candidate genes need to be further analyzed at protein levels.

In conclusion, our study analyzed the transcriptional landscape of human arrested embryos at different stages and identified candidate gene sets and pathways that differed between arrested and developed embryos. Although no DEGs were identified between low- and high-quality late blastocysts, considerable DEGs were identified between day 5 and day 6 blastocysts. Embryo development associated with DNA methylation and piRNA metabolic pathway might be a new filed to investigate. Thus, our study identifies novel markers for embryonic development potential which might aid the choice of blastocysts to be transferred.

## Materials and Methods

### Ethics approval and subjects

This study was approved by the Ethics Committee of Human Study at the Sun Yat-sen Memorial Hospital of Sun Yat-sen University, and the principles of the Declaration of Helsinki were followed. Sixty embryos and 21 couples who underwent PGT between January 2019 and December 2019 were included in this study (Supplemental Table 1). Patients without available blastocysts were excluded from this study. All patients underwent genetic counseling and signed a consent form approved by the local ethics committee. All embryos used in this study were donated and stored for research purposes at Sun Yat-sen Memorial Hospital of Sun Yat-sen University.

### Embryo culture and evaluation

The oocytes were assessed for fertilization approximately 20 h post-injection by checking for the presence of two pronuclei and the second polar body. Embryos were cultured in G1-plus/G2-plus (Vitrolife, Sweden) sequential media in a humidified atmosphere containing 5% O_2_ and 6% CO_2_. According to consensus scoring system for cleavage-stage embryos, blastomere stage embryos on day 3 with less than 10% fragmentation, stage-specific cell size, and no multinucleation were considered grade 1, and blastomere stage embryos on day 3 with 10–25% fragmentation, stage-specific cell size for majority of cells, and no multinucleation were considered grade 2 (Alpha Scientists in Reproductive Medicine and ESHRE Special Interest Group of Embryology. 2011). According to the Gardner grading system, blastocysts graded as 3–6, with either ICM or TE graded above C, were considered suitable late blastocysts for biopsy. Blastocysts graded as 1–2 were considered early blastocysts (Gardner et al. 1999). Based on their scores, the late blastocysts were divided int three categories: ICM and TE ≥ BB (high-quality late blastocysts), ICM and TE < BB but > CC (low-quality late blastocysts), and ICM and TE = CC (unavailable late blastocysts) (Fig. 1A). Available late blastocysts, including low- and high-quality blastocysts, can be used for biopsy. TE biopsy was performed on day 5 (B5) or day 6 (B6) by zona pellucida drilling with a laser, and a few biopsied TE cells (5−8 cells) were removed and collected for genetic analysis (Harton et al. 2011). A single-arrest embryo or chromosomal abnormal blastocyst diagnosed following PGT was vitrified in sequence after the biopsy following the manufacturer’s protocols (Kitazato Corporation, Japan).

### Embryo collection and whole-transcriptome amplification

To remove the zona pellucida, the embryos were exposed to acidic Tyrode’s solution (pH 2.5, Sigma, Germany) for 3–5 s and then washed thoroughly in phosphate-buffered saline (PBS) (pH 7.4, Gibco, USA). Single zona pellucida-free embryos were carefully pipetted and placed into individual tubes containing 2 μL PBS. Whole-transcriptome amplification (WTA) and cDNA generation of whole embryos were performed using the multiple displacement amplification (MDA) reaction (REPLI-g WTA Single Cell, Qiagen, Germany), according to the manufacturer’s protocol.

### RNA-seq library preparation, sequencing-quality control, and expression quantification

All MDA products were used for library construction using the TruePrep DNA Library Prep Kit v2 with MGI adapters (Vazyme, China) according to the manufacturer’s instructions. After the libraries were amplified with phi29 to make DNA nanoballs (DNBs), we used an Agilent Technologies 2100 bioanalyzer (Agilent Technologies, Germany) to assess the quality of the libraries. The libraries were used for RNA-seq performed with a MGISEQ-2000 sequencer (MGI, China). The sequencing procedure yielded bidirectional sequencing with read lengths of 100 bp. The sequencing data were filtered using SOAPnuke software (v1.5.2) (https://github.com/BGI-flexlab/SOAPnuke) (Li et al. 2008). Clean reads were mapped to the reference genome (GRCh38.p12) using HISAT2 (v2.0.4) (http://www.ccb.jhu.edu/software/hisat/index.shtml) (Kim et al. 2015). Bowtie2 (v2.2.5) (http://bowtiebio.sourceforge.net/%20Bowtie2%20/index.shtml) (Langmead et al. 2012) was used to align the clean reads to RefSeq transcripts for quantification. Gene expression levels were quantified using the transcripts per kilobase of exon model per million mapped reads (TPM) using RSEM (v1.2.12) (https://github.com/deweylab/RSEM) (Li et al. 2011).

### Differential expression, enrichment, and Circos analyses

Differential expression analysis was performed using DESeq2 (v1.4.5) (http://www.bioconductor.org/packages/release/bioc/html/DESeq2.html) (Love et al. 2014). We defined upregulated genes (fold change (FC) > 2 and *Q* < 0.05) and downregulated genes (FC < 0.5, *Q* < 0.05) as DEGs between two groups. GO (http://www.geneontology.org/) enrichment analysis of annotated DEGs was performed based on a hypergeometric test (https://en.wikipedia.org/wiki/Hypergeometric_distribution). The GO bar, bubble, and volcano plots were generated using the ggplot2 R package (https://cran.r-project.org/web/packages/ggplot2/index.html). The GO categories in the biological networks were performed using the Cytoscape plug-in ClueGo (https://apps.cytoscape.org/apps/cluego). Heatmaps with H-clusters were drawn using the pheatmap R package (v1.0.8) (https://cran.r-project.org/web/packages/pheatmap/index.html) and TBtools (Chen et al. 2020). Circos analysis was performed using OmicStudio tools at https://www.omicstudio.cn/tool/ (Gu et al. 2014). Ericscript (http://ericscript.sourceforge.net/) was used for gene fusion analysis. Junction reference strictly depends on the Ensembl transcriptome. All reads were mapped against the exon junction reference, and gene fusion products with an EricScore > 0.5 were reported. The 100 bp window sequence map ≥80% of its length against one of the two candidate fused genes was removed (Benelli et al. 2012).

### Blastocyst age and clinical pregnancy rate calculation

The effects of gene expression on the available late blastocyst age were estimated using the following linear model:

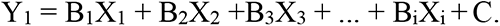

Here, we define the parameter set “B6 = 0, B5 = 1” as the dependent variable. X represents the gene expression level, B is the coefficient, and C is a constant. The score (Y_1_) was used to assess blastocyst age on day 5 or day 6. DEGs between groups B6 and B5 were assessed as independent variables introduced in the regression equation for the blastocyst age score and eliminated when appropriate. Predictors of blastocyst age score were assessed using forward stepwise linear regression analysis, and R^2^ was used to evaluate the goodness of fit of the model. The threshold for statistical significance was *P* < 0.05 (Supplemental Table 2).

The effects of gene expression on the CPRs of available late blastocysts with different ICM scores were estimated using the following linear model:

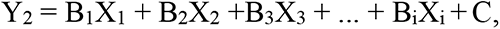

where Y_2_ is the CPR of available late blastocysts with different ICM grades and X is the gene expression level. According to the respective data of single blastocyst transfers performed in our hospital from January 2011 to December 2018, 380 blastocysts were transferred on day 5 and 151 blastocysts on day 6. The CPRs of B5 and B6 were 45.5% (173/380) and 21.9% (33/151), respectively. The CPRs of these five groups were 50.0% (20/40), 44.9% (105/234), 42.4% (28/66), 33.3% (2/6), 22.2% (22/99), 19.6% (9/46), respectively. Predictors of the CPR score were assessed using forward stepwise linear regression analysis, and R^2^ was used to evaluate the goodness of fit of the model. The threshold for statistical significance was *P* < 0.05 (Supplemental Table 3).

### Statistical analysis

All statistical analyses were conducted using SPSS version 25.0 (IBM, USA). Continuous data are presented as mean ± SD for normally distributed variables and as median ± quartile for non-normally distributed variables. Categorical variables were summarized as frequencies and percentages. Comparisons between groups were performed using the chi-square test or Fisher’s exact test for categorical variables. The Kruskal–Wallis and Mann–Whitney U tests were used to assess the difference between non-normally distributed continuous variables. All statistical tests were two-tailed, and a *P-*values < 0.05 were considered statistically significant.

## Data availability

The data pertaining to this article will be shared upon reasonable request to the corresponding author, and the accession number will be disclosed after acceptance.

## Acknowledgments

The authors thank all patients for their participation in this study. This work was partially supported by the National Natural Science Foundation of China (No. 81801431 to P. Y., 81872295 to Y. G. and 32170641 to Y. G.), the Chinese Medical Association clinical medical research special fund,Research and development of young physicians in reproductive medicine (18010060735 to P. Y.); Guangdong Science and Technology Department (2021A1515011183 to Y. G., 2020B1212060018 to Y. G., 2020B1212030004 to Y. G., 2019A1515012005 to W. W. and 2022A1515011152 to W. W.).

## Conflicts of interest

All authors declare that they have no conflicts of interest.

## Supplemental Figure legends

Supplemental Fig. 1 Reference genome alignment and expression distribution

A) Mapping ratio of genome.

B)Mapping ratio of RefSeq transcripts.

C)Mapping ratio of mitochondria

D)Stacked column chart of expression distribution.

Supplemental Fig. 2 Differential gene expression (DEG) analysis among different groups.

A)Number of upregulated DEGs and downregulated DEGs when comparing samples from adjacent stages.

B)Volcano chart of DEGs between 4C_AB vs 8C_AB.

C)Expression of the six DEGs from supplementary Fig. 3B.

Supplemental Fig. 3 Expression heatmap of gene modules related to cell fate transcriptional signatures.

Expression heatmap of gene modules related to cell fate transcriptional signatures in normally developing human embryos. The row represents the gene and the column represents the median expression level of a developmental stage. The red fonts indicate the genes with a significant difference (*P* < 0.025, single-tail test) when comparing the expression levels of samples on either side of the crack.

Supplemental Fig. 4 Differential gene expression (DEG) analysis between B5s and B6s. Volcano chart of DEGs between B6s vs B5s.

Supplemental Fig. 5

Expression level of four maternal genes analyzed in our study (A-D) and in the study of Yan et al. (E-H). * *P* < 0.05; **** *P* < 0.0001.

## Supplemental Tables 1-6

Supplemental Table 1. Clinical features of 60 embryos from 21 couples that underwent preimplantation genetic testing.

Supplemental Table 2. Linear regression equation of the gene expression effects on blastocyst age.

Supplemental Table 3. Linear regression equation evaluated of the gene expression effects on clinical pregnancy rates of blastocysts with different inner cell mass (ICM) scores.

Supplemental Table 4. Quality control of RNA-seq data from 60 embryos.

Supplemental Table 5. Raw transcripts per kilobase of exon model per million mapped reads (TPM) gene expression data of 60 embryos.

Supplemental Table 6. Comparison of blastocyst and available late blastocyst rates between PGT-SR and IVF groups; PGT-SR group: preimplantation genetic testing for chromosomal structural rearrangement in couples); IVF group: traditional IVF couples with normal chromosol karyotypes.

## Notes

### Competing Interest Statement

The authors have declared no competing interest.

